# KRAS G12D can be targeted by potent salt-bridge forming inhibitors

**DOI:** 10.1101/2021.12.13.472365

**Authors:** Zhongwei Mao, Hongying Xiao, Panpan Shen, Yu Yang, Jing Xue, Yunyun Yang, Yanguo Shang, Lilan Zhang, Xin Li, Yuying Zhang, Yanan Du, Chun-Chi Chen, Rey-Ting Guo, Yonghui Zhang

## Abstract

KRAS mutation occurs in nearly 30% of human cancers, yet the most prevalent and oncogenic KRAS mutation (G12D) still lacks inhibitors. Herein, we explored the formation of a salt-bridge between KRAS’s Asp12 residue and a series of potent inhibitors. Our ITC results show that these inhibitors bind to and inhibit both GDP-bound and GTP-bound KRAS G12D, and our crystallographic studies revealed the structural basis of inhibitor binding in the switch-II pocket, experimentally confirming the formation of a salt-bridge between the piperazine moiety of the inhibitors and the 12D residue of the mutant protein. Among KRAS family proteins and mutants, both ITC and enzymatic assays support the selectivity of the inhibitors for KRAS G12D, and the inhibitors disrupt the KRAS-CRAF interaction. We also observed inhibition of cancer cell proliferation and inhibition of MAPK signaling by a representative inhibitor (TH-Z835); however, since this was not fully dependent on KRAS mutation status, it is possible that our inhibitors may have off-target effects via non-KRAS small GTPases. Experiments with a mouse model of pancreatic cancer showed that TH-Z835 significantly reduced tumor volume and synergized with an anti-PD-1 antibody. Collectively, our study demonstrates proof-of-concept for a salt-bridge, induced-fit pocket strategy for KRAS G12D, which warrants future medicinal chemistry efforts for optimal efficacy and minimized off-target effects.

## Introduction

The oncogenic impacts of the *KRAS* gene were first reported in 1980s, making *KRAS* one of the first identified oncogenes^1^. And it is now understood that the KRAS protein functions as a molecular switch controlling multiple signaling pathways, which respond to upstream EGFR activation and regulate the downstream MAPK and PI3K/mTOR pathways, eventually instructing cell proliferation, differentiation, and survival^2,3^.

Clinical data has implicated driver mutations of the KRAS residue G12, and basic studies have shown that such mutations impair both this enzyme’s intrinsic and GAP (GTPase-activating protein)-stimulated GTP hydrolysis activity^7^, which promotes oncogenesis. Despite nearly four decades of efforts, no direct KRAS inhibitor has been approved for medical use. There is consensus that the difficulty in developing direct KRAS inhibitors relates on the one hand to the picomolar affinity of GTP and GDP to KRAS (and the high intracellular concentrations of these metabolites), and on the other hand to an absence of suitable deep pockets for allosteric regulation.

One major breakthrough for KRAS inhibition was the discovery of an allosteric switch-II pocket (S-IIP) that is induced by covalent inhibitors of KRAS bearing the G12C driver mutation^8^. Studies have shown that induction of S-IIP results from covalent bond formation between the electrophilic acryloyl moieties of these inhibitors and the nucleophilic thiol moiety of the Cys residue at position 12^9–16^. These KRAS G12C inhibitors have shown promising results in recent clinical trials, although it is notable that they exclusively target inactive state KRAS^9^. Despite these developments with inhibitors of KRAS G12C, the most prevalent and oncogenic G12 mutant variant is KRAS G12D, which is estimated to impact more than 50% patients of pancreatic ductal adenocarcinoma^17^.

Recent studies have explored various approaches for targeting KRAS G12D, including indole-based small molecules targeting a switch-I/II pocket, a compound (KAL-21404358) targeting the P110 site, use of a pan-RAS inhibitor (compound 3144) targeting the A59 site, and a cyclic peptide (KD2) targeting KRAS G12D^18–21^. However, none of these molecules target KRAS G12D with suitably low (micromolar) activity. The genesis of the present study was our speculation that a compound targeting the aspartic acid residue of KRAS G12D that could somehow mimic the covalent bond *(i.e.,* that formed between inhibitors and the cysteine residue of KRAS G12C) may likewise induce an allosteric pocket. This would perhaps enable pharmacological inhibition of this more prevalent oncodriver-mutation-bearing KRAS variant.

## Results

### A salt-bridge based strategy for targeting KRAS G12D with a methyl-substituted piperazine inhibitor

Assuming that the α-carboxylic acid moiety of Asp12 is deprotonated under physiological conditions, we pursued a strategy of forming a strong interaction targeting Asp12 based on generating a salt-bridge interaction with an alkyl amine moiety on an inhibitor. As the overall structures of the G12C and KRAS G12D variants are highly similar, we started our efforts based on a scaffold of the G12C inhibitor MRTX^22^ (**Fig. S1a**). Removal of MRTX’s acryloyl moiety exposed a piperazine moiety that was predicted to position this alkyl amine in close enough proximity to Asp12 (2.2 Å) to support salt-bridge interaction (**Fig. S1b**).

Pursuing this, we synthesized TH-Z801 (**Fig. 1a**), which exerted relatively weak inhibition (IC_50_ 115 μM), assessed based on the GDP/GDP exchange rate of KRAS G12D as catalyzed by SOS^9^. Additional SAR studies showed that replacement of the piperazine moiety with non-amine moieties dramatically decreased the inhibitory activity, supporting the functional contribution of a salt-bridge interaction in slowing down the GDP/GDP exchange rate (**Fig. 1a, Supplementary Table 1**).

**Fig. 1:**
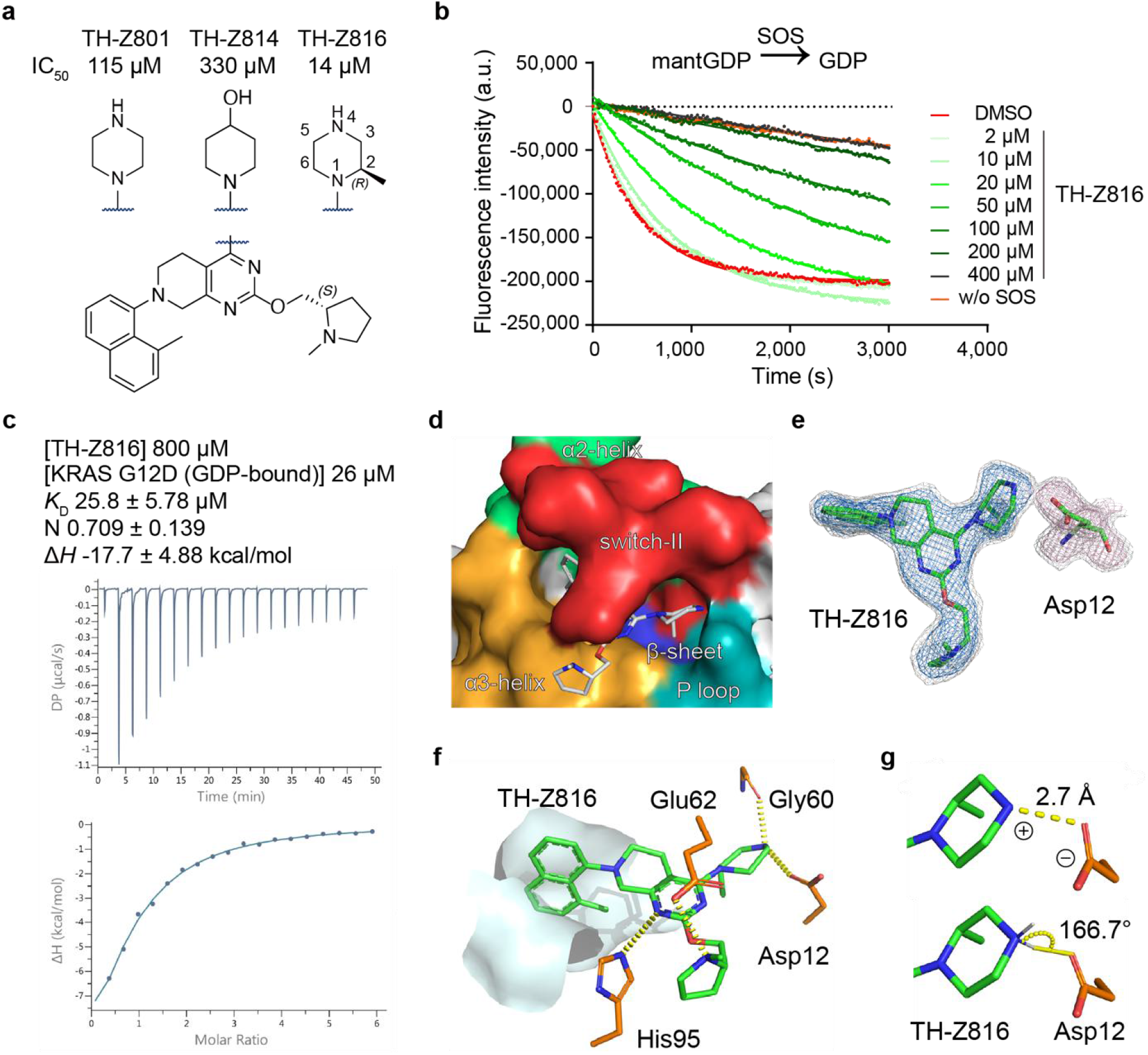
A piperazine-focused strategy for targeting KRAS G12D with salt-bridge interaction. **a**, Chemical structures of TH-Z801and TH-Z816 with exposed piperazines, structure of TH-Z814 with non-amine moiety. **b**, Inhibitory activities of TH-Z816 on KRAS G12D measured by SOS-catalyzed nucleotide exchange assay with GDP as the incoming nucleotide. **c**, ITC assay of the TH-Z816 (800 μM) and GDP-bound KRAS G12D (26 μM). **d**, Crystal structure (PDB ID 7EW9) of KRAS G12D bound to TH-Z816 (white stick). The binding pocket is formed by the α2-helix (green), switch-II (red), α3-helix (orange), P loop (teal), and the central β-sheet (blue) of KRAS G12D. **e**, *Fo-Fc* maps of TH-Z816 (blue mesh, 2.5 σ; gray mesh, 1.5 σ) and Asp12 (pink mesh, 2.5 σ; gray mesh, 1.5 σ). **f**, Interactions between TH-Z816 (green) and surrounding residues (orange). The hydrophobic sub-pocket is shown as a surface diagram. **g**, The ionic bond (upper panel) and hydrogen bond (lower panel) between piperazine and Asp12.

Further chemical exploration focusing on piperazine substitution yielded TH-Z816 (wherein the piperazine was (*R*)-methyl-substituted), which had relatively strong inhibition activity (IC_50_ 14 μM) (**Fig. 1a, 1b**). We further conducted isothermal titration calorimetry (ITC) assays to test whether TH-Z816 can bind directly to KRAS G12D: indeed, the detected binding affinity (K_D_) of TH-Z816 with KRAS G12D (GDP-bound) was 25.8 μM (**Fig. 1c**).

### A complex structure revealed an induced-fit pocket and confirmed the salt-bridge interaction

We solved a 2.13 Å co-crystal structure of KRAS G12D in complex with TH-Z816 (**Table S1**). Our structure showed that TH-Z816 induced an allosteric pocket positioned under the KRAS G12D switch-II region (**Fig. 1d**); no such pocket was evident in the inhibitor-free structure (**Fig. S1c**). This induced-fit pocket was located near the α2-helix, switch-II, α3-helix, the P loop, and the central β-sheet of KRAS (**Fig. 1d**). Note that the pocket shape was quite similar (RMSD 0.293 Å, 148 to 148 atoms) to the MRTX-induced SII-P of KRAS G12C (**Fig. S1d**).

Well-defined electron density clearly supported the conformation of TH-Z816 and KRAS G12D (**Fig. 1e**). Specifically, the naphthyl moiety of TH-Z816 embedded deeply into the pocket and formed hydrophobic interactions with residue Val9, Met72, Phe78, Gln99, Ile100, and Val103 (**Fig. S1e**). There are also four pairs of polar interactions between TH-Z816 and residues His95, Glu62, Gly60, and Asp12 (**Fig. 1f**). Note that an ionic bond and a hydrogen bond together comprise the anticipated salt-bridge: on one hand, the positively charged piperazine moiety of TH-Z816 and negatively charged side chain of Asp12 are close enough to support an ionic bond (**Fig. 1g**); on the other hand, the measured bond angle (166.7°) and length (2.7 Å) supports the presence of a hydrogen bond between these two moieties (**Fig. 1g**).

### A cyclization strategy to improve inhibitory potency by optimizing ΔS

Further optimization of TH-Z816 was guided by the following thermodynamic rule, ΔG = ΔH - TΔS^23, 24^. We sought to optimize for ΔS by restraining the conformational freedom of the piperazine moiety. Given the axial position of the methyl group, we employed a cyclization strategy based on TH-Z816 and generated the bicyclic compound TH-Z827 (**Fig. S2a**). As TH-Z827 conferred a more than 8-fold increase in potency (IC_50_ 3.1 μM), we subsequently adopted a ΔS-focused cyclization strategy and designed additional bicyclic compounds (**Fig. 2a, Supplementary Fig. 1**), among which the most potent one was TH-Z835 (IC_50_1.6 μM) (**Figs. 2b, S2b, Supplementary Table 1**).

**Fig. 2:**
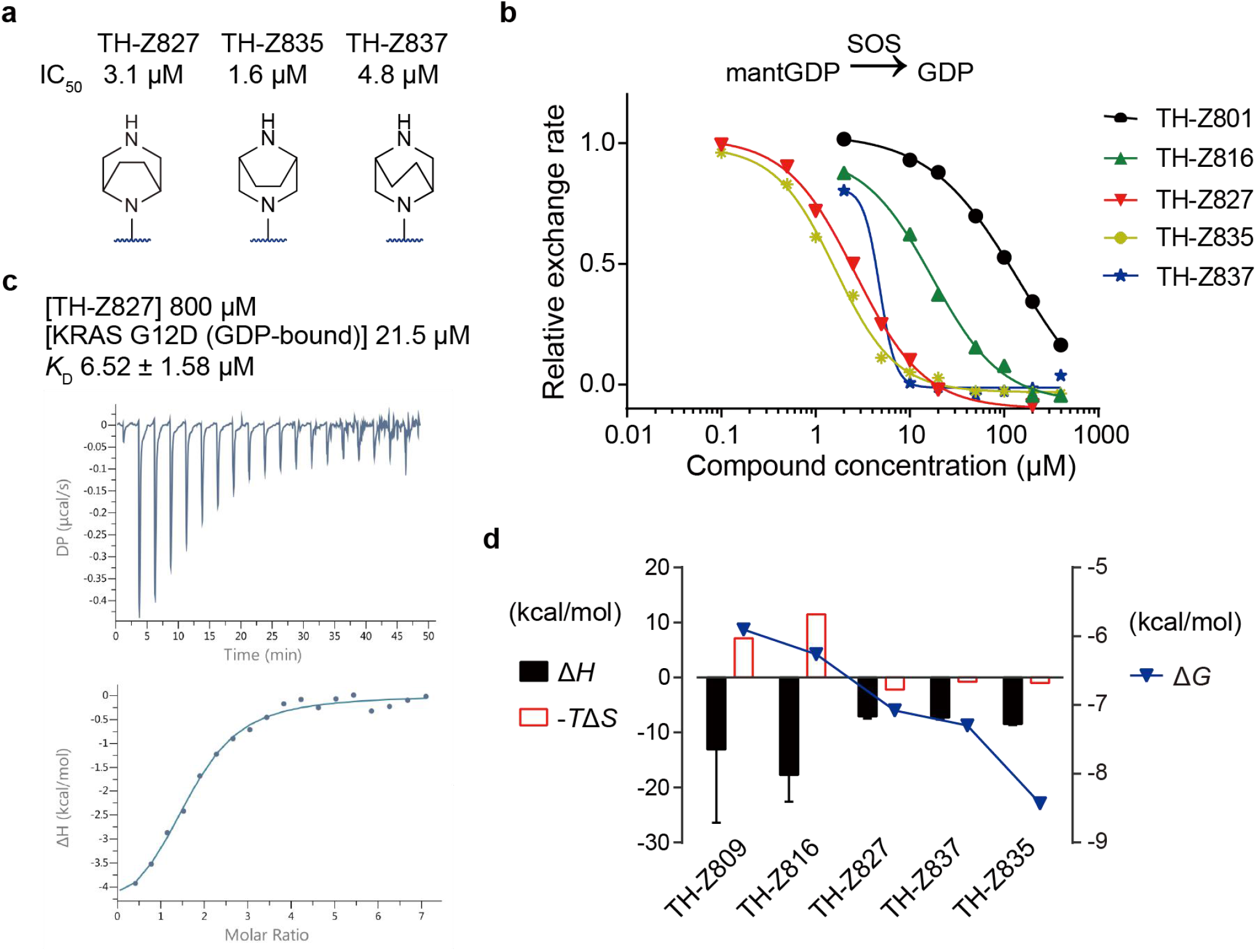
A cyclization strategy to improve inhibitory activities and binding potency. **a**, Chemical structures of bicyclic compounds TH-Z827, TH-Z835, and TH-Z837. **b**, Inhibitory activities of these bicyclic compounds on KRAS G12D measured by SOS-catalyzed nucleotide exchange assay with GDP as the incoming nucleotide. **c**, ITC assay of the TH-Z827 (800 μM) and GDP-bound KRAS G12D (21.5 μM). **d**, ΔG, ΔH and ΔS analysis of compounds tested by ITC assays. For each compound, ΔH and −TΔS values are presented at the left axis, while the ΔG value is presented at the right axis.

We next conducted ITC assays to measure the binding affinity of these bicyclic compounds to KRAS G12D in the presence of GDP (**Fig. 2c, S2c**). Compared with TH-Z816, bicyclic compounds TH-Z827, TH-Z835, and TH-Z837 had unfavorable ΔH changes yet favorable ΔS changes (**Fig. 2d**), indicating that the binding affinity (ΔG) increase of these bicyclic compounds can be attributed to ΔS improvement.

### Co-crystal structures showed G12D inhibitors bind to GDP-bound and GMPPNP-bound KRAS

Previous studies have shown that KRAS exists in cells in both GTP-bound and GDP-bound states^13^. A molecular docking analysis of our KRAS G12D inhibitors indicated that the absence of an acryloyl moiety in the inhibitors should leave sufficient space (4.7 Å) for the γ-phosphate of GTP, while maintaining the salt-bridge interaction between Asp12 of KRAS and the piperazine moiety of our inhibitors (**Fig. S3a**).

To reveal whether these inhibitors bind to the GTP-bound KRAS G12D, we first treated the purified protein with GMPPNP (a stable analog of GTP) and EDTA, following a well-established nucleotide-exchange protocol^8,9^. We next initiated a systematic effort aimed at crystallizing GMPPNP-containing KRAS G12D in complex with these more active inhibitors TH-Z827 and TH-Z835, and successfully solved a 2.25 Å co-crystal structure of KRAS G12D in complex with TH-Z827 (PDB ID 7EWA, **Fig. 3a, Table S1**). This crystal structure was trimeric, with monomer A bound to GDP and TH-Z827, monomer B bound to GMPPNP and TH-Z827, and monomer C bound to GMPPNP. We also solved a co-crystal structure of KRAS G12D in complex with TH-Z835 (PDB ID 7EWB, **Fig. 3a, Table S1**). In these two inhibitor-complex structures, the electron densities were well-defined for inhibitors, for Asp12, and for both GMPPNP and GDP (**Fig. 3a**), and our data confirmed that both TH-Z827 and TH-Z835 are able to interact with Asp12 and form a salt bridge for both GDP-bound and GMPPNP-bound KRAS G12D. It is noted that in the inhibitor-free GMPPNP-bound state, the conformation of switch II is more tightly bound to the γ-phosphate. However, binding of TH-Z835 shifted the conformation of switch II towards the inhibitor-free GDP-bound conformation (analysis shown in **Supplementary Figure 2**).

**Fig. 3:**
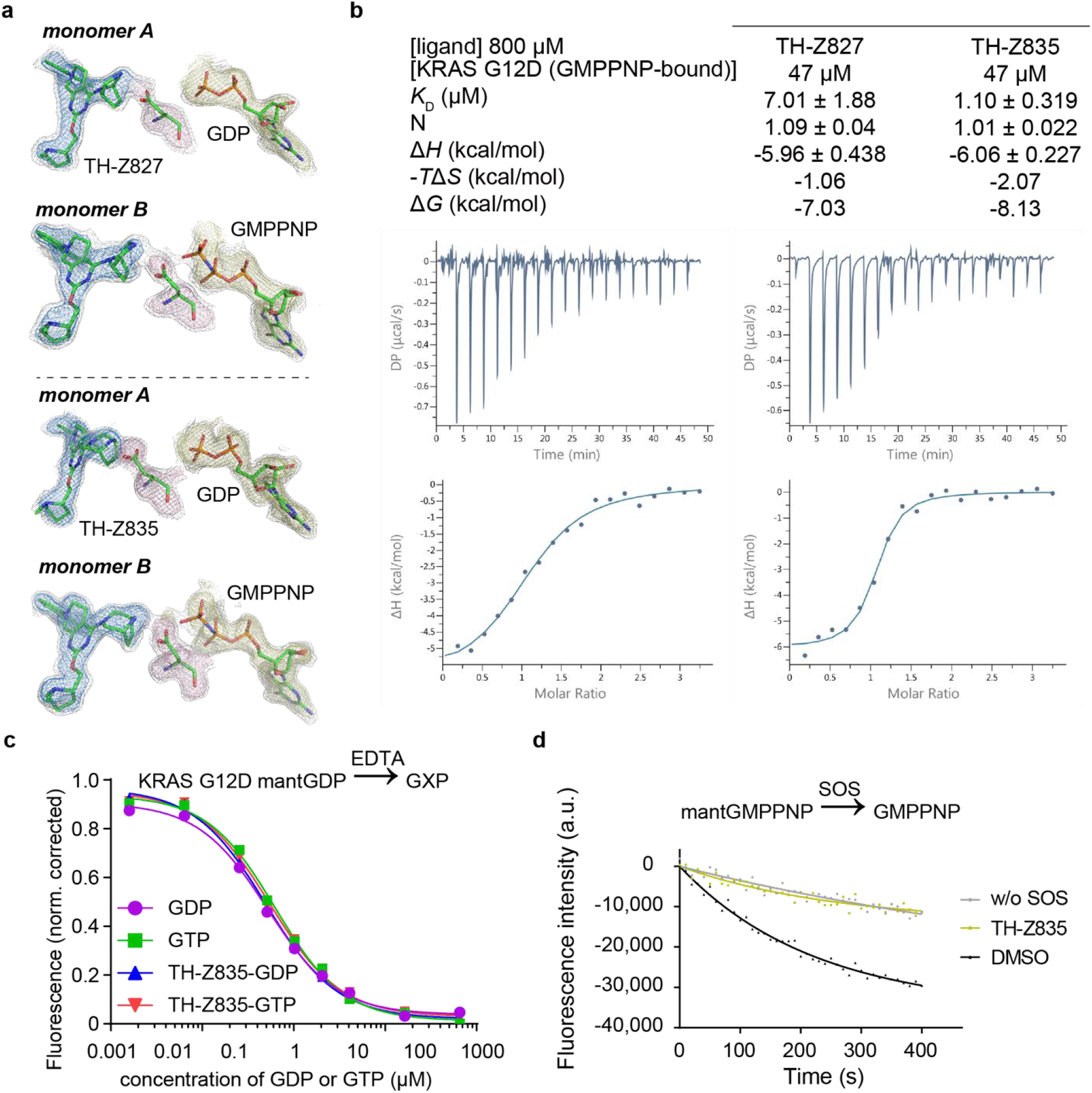
G12D inhibitors bind to both GDP-bound and GMPPNP-bound KRAS. **a**, (upper panel) *Fo-Fc* map of TH-Z827, Asp12, and GDP (PDB ID 7EWA, monomer A). *Fo-Fc* map of TH-Z827, Asp12, and the GTP analog GMPPNP (PDB ID 7EWA, monomer B). (lower panel) *Fo-Fc* map of TH-Z835, Asp12, and GDP (PDB ID 7EWB, monomer A). *Fo-Fc* map of TH-Z835, Asp12, and GMPPNP (PDB ID 7EWB, monomer B). In all graphs, the 1.5 σ *Fo-Fc* maps of all shown elements are shown in gray mesh. The color scheme of 2.5 σ *Fo-Fc* maps is: blue for inhibitors, pink for Asp12, and yellow for nucleotides. **b**, ITC assay of the TH-Z827 or TH-Z835 (800 μM) with GMPPNP-bound KRAS G12D (47 μM). **c**, EDTA-mediated competition between fluorescently labeled mantGDP loaded on KRAS and free nucleotide (GDP or GTP). The experiment was carried out with KRAS G12D alone or with KRAS G12D treated with TH-Z835. **d**, Inhibitory activity of TH-Z835 measured by SOS-catalyzed nucleotide exchange assays with GMPPNP as the incoming nucleotide

### The inhibitors have similar binding affinities for GDP-bound and GMPPNP-bound KRAS

We next conducted ITC assays to measure the binding affinities between G12D inhibitors and GMPPNP-bound KRAS G12D (**Fig. 3b, S3c**). The detected binding affinities for each of the tested compounds were within a narrow range for both the GDP-bound and the GMPPNP-bound forms (**Fig. S3b**). And EDTA-catalyzed nucleotide exchange assays to test the GDP/GTP binding preference of KRAS G12D supported our ITC results, showing no GDP/GTP binding preference for KRAS G12D in the presence of our inhibitors (**Fig. 3c**). In contrast, KRAS G12C had significantly lower affinity for GTP than for GDP in the presence of the G12C inhibitor MRTX (**Fig. S3d**), which was consistent with previous reports^9^. We also conducted SOS-catalyzed nucleotide exchange assays of KRAS G12D, which experimentally confirmed that our inhibitor TH-Z835 does inhibit both mantGMPPNP/GPPNP exchange and GPPNP/mantGMPPNP exchange (**Figs. 3d, S3e**).

### Mutant-selectivity of the KRAS G12D inhibitor TH-Z827

We next studied the mutant-selectivity of our G12D inhibitors using ITC assays: no measurable binding was detected for TH-Z827 with KRAS WT or with KRAS G12C, no matter whether the proteins were GDP-bound or GMPPNP-bound **(Fig. 4a, Fig. S4**). SOS-catalyzed nucleotide exchange assays also showed that TH-Z827 exerted a nearly 10-fold stronger inhibition for KRAS G12D than for KRAS G12C (IC_50_ 2.4 μM vs 20 μM) (**Fig. 4b**).

**Fig. 4:**
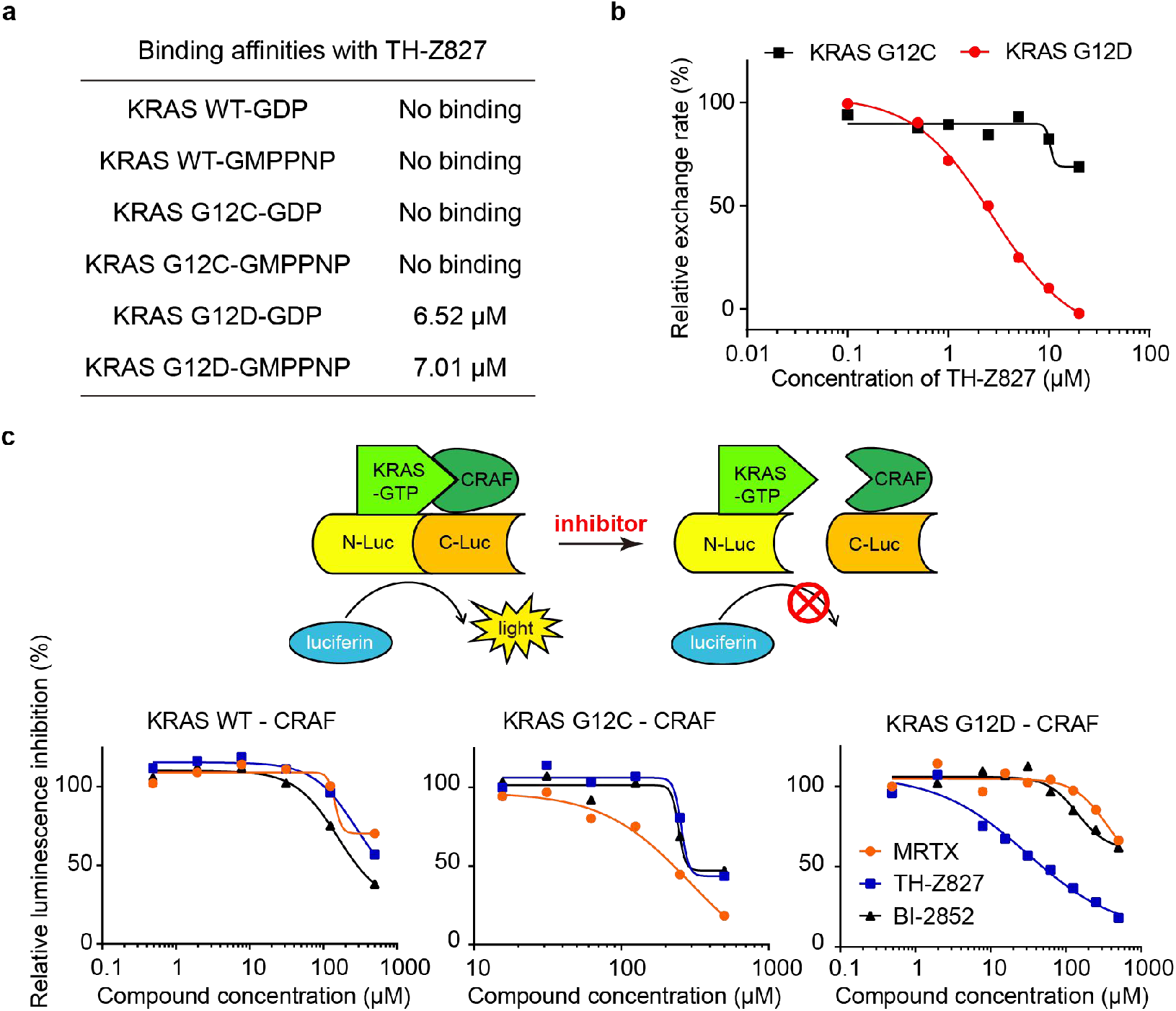
Mutant-selectivity of the KRAS G12D inhibitor TH-Z827. **a**, Binding affinities between TH-Z827 and GDP- or GMPPNP-bound KRAS (WT, G12C, or G12D). **b**, SOS-catalyzed KRAS G12C or G12D nucleotide exchange assays in the presence of TH-Z827. **c**, Principle of the split-luciferase reporter assays detecting the KRAS-CRAF interaction in lysates from HEK 293T cells, with or without TH-Z827. Two other compounds (the G12C inhibitor MRTX and the pan-RAS inhibitor BI-2852) were included as controls.

Confirming the mutant-selectivity of targeting KRAS alone, we further studied whether our inhibitor could selectively block the interactions of various KRAS mutants with its effector protein. CRAF is a well-studied KRAS effector protein, and the KRAS-CRAF interaction is known to promote MAPK signal transduction^31–33^. We established a split-luciferase reporter system based on HEK 293T cells (**Fig. 4c**). Upon doxycycline treatment, the cells express both full-length KRAS (fused to N-luciferase) and a truncated CRAF variant (comprising residue 50–220) fused to C-luciferase. After lysis, the supernatant was incubated with our inhibitor and the substrate luciferin. When KRAS binds to CRAF, luciferase complementation results in emission of a luminescence signal. The results showed that TH-Z827 blocks the KRAS G12D-CRAF interaction with an IC_50_ 42 μM but does not affect CRAF’s interaction with KRAS WT or KRAS G12C at this concentration (**Fig. 4c**).

### Anti-proliferative effects and MAPK signaling inhibition

The promising inhibitory activity and mutant-selectivity results from the ITC and KRAS-CRAF interaction assays motivated us to evaluate the potential anti-cancer effects of TH-Z827. The KRAS G12D mutation is the most prevalent driver gene for the most prevalent form of pancreatic cancer^34^. In two pancreatic cancer cell lines bearing the KRAS G12D mutation (PANC-1 and Panc 04.03), TH-Z827 conferred anti-proliferative effects with IC_50_ values of 4.4 μM and 4.7 μM, respectively (**Fig. S5a**). Treatment with TH-Z827 also reduced the levels of pERK and pAKT in PANC-1 and Panc 04.03 cells (**Fig. S5b**), confirming that TH-Z827 prevented the activation of MAPK and PI3K/mTOR signaling. We observed that TH-Z835 reduced the pERK level in PANC-1 cells with an IC_50_ less than 2.5 μM (**Fig. 5c**), which was more potent than TH-Z827 (**Fig. S5b**). These findings are consistent with their binding affinities as measured in the ITC assays (**Fig. 3b**) and their anti-proliferative IC_50_ values (**Fig. S5a**). We next performed 2D adherent, 3D non-adherent, and colony formation assays and observed anti-proliferative effects from TH-Z835 for two KRAS G12D-bearing pancreatic cancer cell lines: PANC-1 and KPC (KrasLSL.G12D/+; p53R172H/+; PdxCretg/+). It was notable that the IC_50_ values of TH-Z835 in the colony formation assay were less than 0.5 μM (**Figs. 5a, 5b**).

**Fig. 5:**
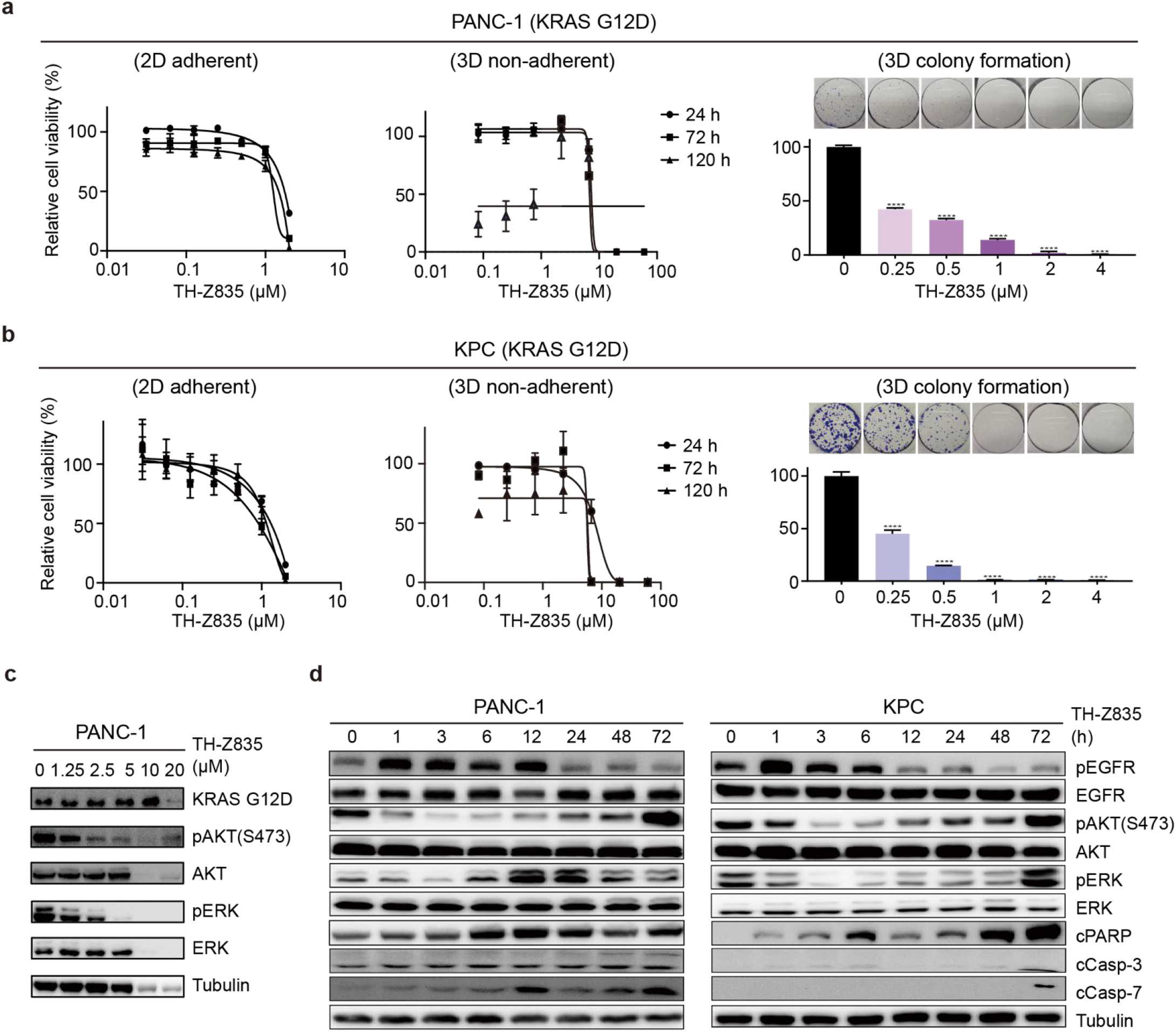
Anti-proliferative effects of TH-Z835. **a-b,** Cell viability assays of PANC-1 and KPC cells treated with indicated concentration of TH-Z835 for 24 h, 72h, or 120 h in 2D adherent assays (**left panel**) and 3D non-adherent assays plates (**middle panel**). As for colony formation assay (**right panel**), PANC-1 cells were cultured for 14 days and KPC cells were cultured for 10 days. **c**, Immunoblotting analysis of ERK and AKT phosphorylation status in PANC-1 cells treated with TH-Z835 for 3 h. **d**, Immunoblotting of EGFR, ERK, AKT phosphorylation, cleaved PARP (cPARP), cleaved Caspase-3 (cCasp-3), and cleaved Caspase-7 (cCap-7) in PANC-1(**left panel**) and KPC (**right panel**) cells treated with TH-Z835 (5 μM).

Further examination of the inhibitor-treated PANC-1 cells revealed increased protein levels for p21 and p27, as well as decreased levels of CDK2/4/6 and Cyclin D1, indicating that TH-Z835 induces arrest at the G1 phase of the cell cycle (**Fig. 6a**). Consistent with these western blotting results, flow cytometry also showed an increased population of PANC-1 cells in G1 phase and a decreased population in S phase as compared with the DMSO control group (**Fig. 6b**). We also used flow cytometry to evaluate apoptosis induction in TH-Z835 treated PANC-1 and KPC cells and found significantly increased proportions of Annexin V positive cells in cultures of KPC cell lines (**Fig. 6c**). We confirmed this finding by immunoblotting which showed that treatment with TH-Z835 led to increased levels of the apoptosis markers including cleaved PARP, cleaved caspase-3, and cleaved caspase-7 (**Figs. 5d, 6a**).

**Fig. 6:**
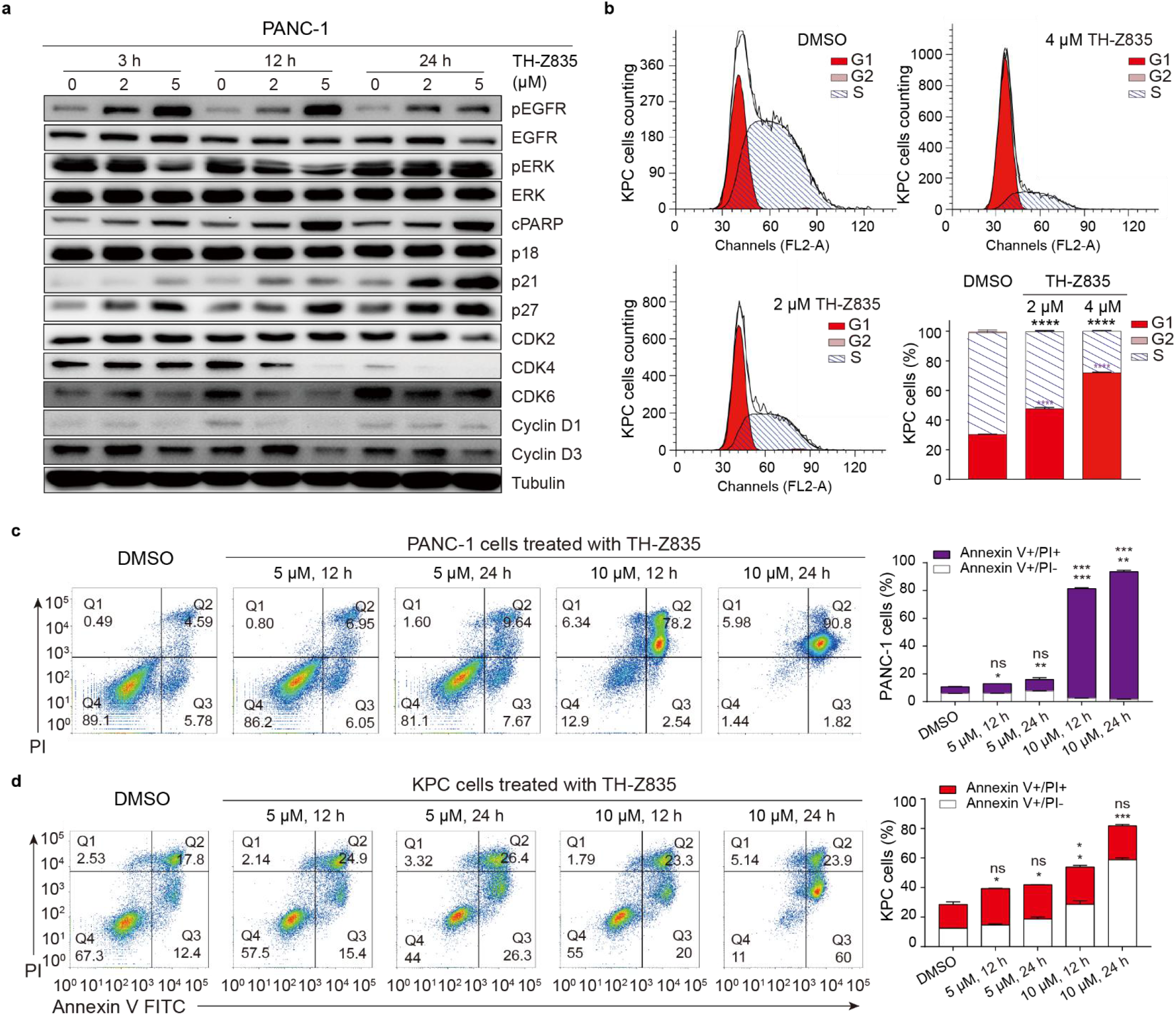
TH-Z835 induced cell cycle arrest and apoptosis in KRAS G12D mutant cells. **a**, Immunoblotting against RTK signaling and cell cycle marker proteins in PANC1 cells after 3-, 12- or 24-h treatment with the indicated concentrations of TH-Z835. **b,** Percentages of KPC cells in the G1, S, and G2 phases after treated with TH-Z835 for 24 h. Data are shown as the mean ± s.e.m. (*n* = 3). Student’s t test, **** p < 0.0001. **c**, **d**, **(left panel)** Cell apoptosis analyzed by flow cytometry of PANC-1 (**c**) or KPC (**d**) cells upon a 12 or 24-h treatment with TH-Z835 (5 μM or 10 μM). **(right panel)** Apoptotic (Annexin V-positive) cell proportions were quantified. Data are show as the mean ± s.e.m. (*n* = 3). Student’s t test, * *p* < 0.05, ** *p* < 0.01, *** *p* < 0.001.

We also tested the antiproliferative effects of TH-Z835 in other non-G12D mutant cancer cell lines, including 4T1 (KRAS WT), MIA PaCa-2 (KRAS G12C), CFPAC-1 (G12V), and HCT116 (G13D) cells (**Fig. S5c**). Against our expectations based on our *in vitro* protein assay data for the KRAS family proteins, these assays indicated that TH-Z835 induced apoptosis (**Fig. S6a-d**) and reduced the pERK and pAKT levels in these cells (**Fig. S5d**). This is not totally unexpected, especially given that proliferation and MAPK signaling induction in diverse cancers has been shown to be regulated by other Ras superfamily proteins, including members of the Rho, Ran, Arf, and Rab families^35^. It should be noted that the switch-II regions of these proteins share structural similarity to KRAS proteins, so it is plausible that these may also be targeted by TH-Z835, which awaits future investigations.

### *In vivo* antitumor activity and combination with PD-1 therapy

Our *in vivo* testing of KRAS G12D inhibitors were conducted in two xenograft pancreatic tumor models: BALB/c nude mice subcutaneously inoculated with Panc 04.03 cells and C57BL/6 mice inoculated with KPC cells. In the nude mice model, TH-Z827 significantly reduced the tumour volumes compared to the vehicle control, and did so in a dose-dependent manner (**Fig. 7a**). However, an intraperitoneal dosing of 30 mg kg^−1^ TH-Z827 resulted in observed weight loss, again suggesting the potential of off-target effects (**Fig. S7a**). We next tested the antitumor activity of TH-Z835 in the C57BL/6 mice model. The tumor volume was also reduced compared with the vehicle group (**Fig. S7d**). And additional immunohistochemical analysis of tumour sections revealed an increased expression of cleaved caspase-3 and a decreased expression of pERK1/2 (**Figs 7c, S7e**), indicating that TH-Z835 induces apoptosis and MAPK signaling inhibition *in vivo.*

**Fig. 7:**
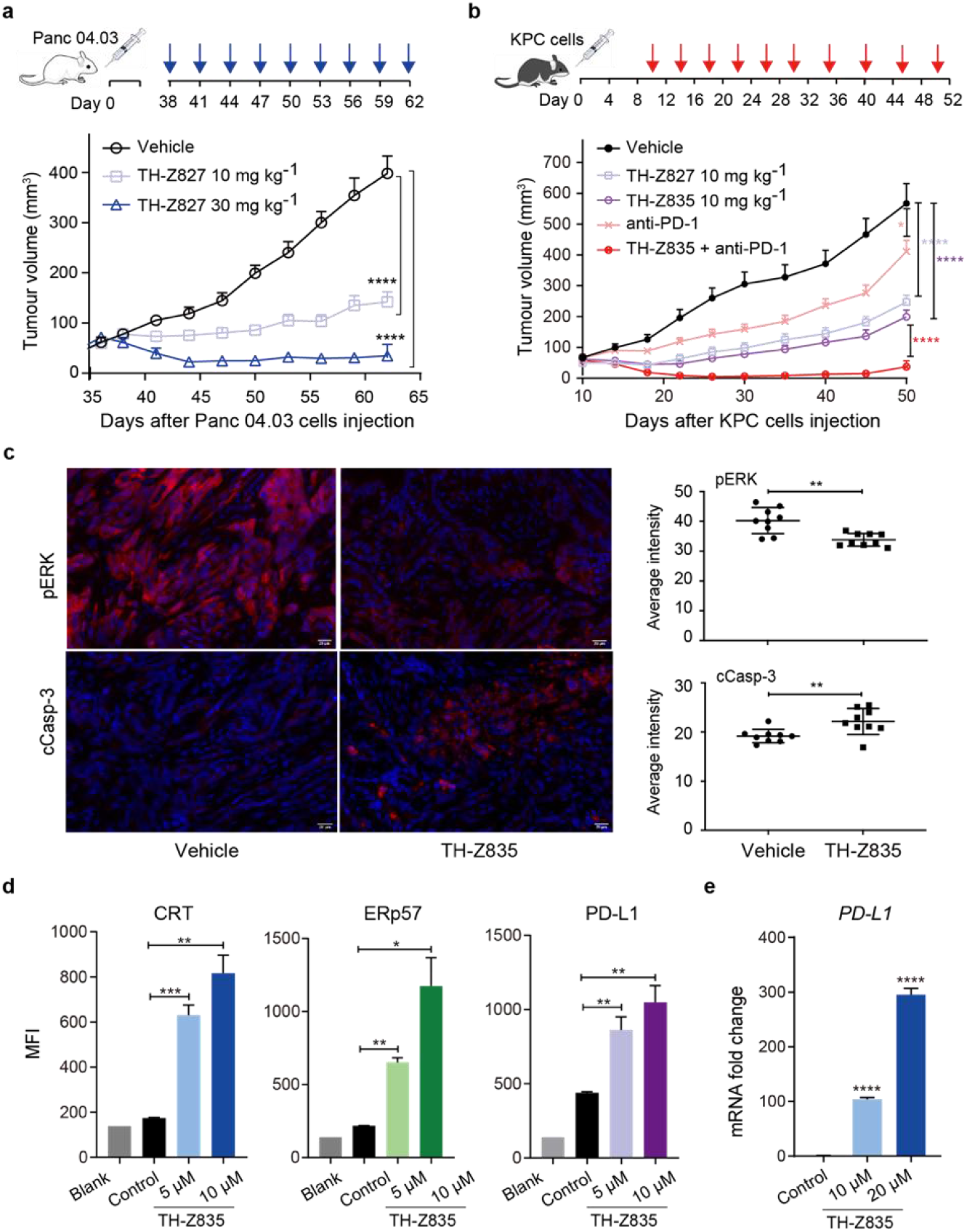
Anti-tumor effects of the KRAS G12D inhibitors alone and in combination with anti-PD-L1 antibody. **a,** Nude mice injected with Panc 04.03 cells at Day 0, were later IP injected with TH-Z827 (at 10 mg kg^−1^ or 30 mg kg^−1^) according to the indicated dosage schedule (blue arrow). The tumour volumes (mean ± s.e.m., *n* = 10) were measured with digital calipers and assessed using one-way ANOVA followed by Dunnett’s test (**** P_adj_< 0.0001). **b,** C57BL/6 mice were injected with KPC cells at Day 0, after which TH-Z827, TH-Z835, anti-PD-1 antibody, or a combination therapy (10 mg kg^−1^ TH-Z835 and 100 μg per dose anti-PD-1 antibody) was IP administered according to the indicated dosage schedule (red arrow). The tumour volumes (mean ± s.e.m., *n* = 5) were assessed using Student’s t test, * *p* < 0.05, ** *p* < 0.01, *** *p* < 0.001) **c,** (**left panel**) Immunofluorescence (IF) analysis of pERK and cleaved Caspase-3 in tumor sections from C57BL/6 mice model (see also **Fig. 7Sd**) treated with TH-Z835 or not. Scale bar, 20 μm. (right panel) quantification of IF positive staining. Data are shown as the mean ± s.d. (*n* = 9), Student’s t test, ** *p* < 0.01. **d,** Flow cytometry analysis of the immunogenic cell death (ICD) markers CRT and ERp57 and the immune checkpoint protein PD-L1 on the surface of KPC cells after 24-h treatment with TH-Z835. **e,** mRNA expression level of *PD-L1* in PANC-1 cells after 24-h treatment with TH-Z835.

Recent studies showed that KRAS oncogenic signaling can extend beyond cancer cells to orchestrate the immune status of the tumour microenvironment^36–38^. Specifically, studies of murine pancreatic cancer models showed that KRAS can drive immune evasion (characterized by scant intratumoural CD8^+^ T cells)^39^. In addition, activated MAPK signaling may also be involved in the immunosuppressive tumor microenvironment^40^. We first tested the effect of TH-Z835 treatment on KPC cells and found increased *PD-L1* mRNA expression levels and increased the surface levels of the immunogenic cell death markers CRT and ERp57 (**Figs. 7d, 7e, S7f**). We next tested the efficacy of a combination therapy comprising TH-Z835 and an anti-PD-1 antibody using the C57BL/6 mice inoculated with KPC cells. Indeed, a combination treatment led to a statistically significant decrease in tumour volume as compared to either of the mono-therapies (**Fig. 7b**). This synergism could also be observed in a combination therapy comprising TH-Z827 and the anti-PD-1 antibody (**Fig. S7c**).

## Discussion

Aiming to develop KRAS G12D inhibitors—specifically using a strategy to mimic the acryloyl-cysteine interaction of KRAS G12C inhibitors with a salt-bridge for the Asp residue of the more common KRAS G12D mutant variant—we used structure-based drug design, synthesized a series of small molecules, and characterized their activities both *in vitro* and *in vivo.* Unlike covalent KRAS G12C inhibitors that selectively bind to GDP-bound proteins, both our computational and ITC investigations suggested that these salt-bridge forming inhibitors can bind to both GDP-bound and GTP bound KRAS G12D. We also solved crystal structures of KRAS G12D with series of potent inhibitors (TH-Z816, TH-Z827, and TH-Z835). Our structural data confirmed that these inhibitors can induce the formation of an allosteric pocket under the KRAS G12D switch-II region, similar to reported findings for the covalent KRAS G12C inhibitors^8, 9, 13, 15^. Secondly, the crystal structures revealed that these inhibitors are able to bind to the GMPPNP-bound KRAS G12D. That is, whereas steric clashing renders the G12C inhibitors incapable of targeting KRAS G12C^9^, our salt-bridge forming inhibitors have sufficient space to target Asp12 in both GDP-bound and GTP-bound KRAS G12D. Notably, the discovery of G12C inhibitors made a stereotype that targeting the GDP-bound (inactive) KRAS may be a more viable approach than targeting the GTP-bound (active) KRAS^5^. Now, using the inhibitors developed in the present study, it should be possible to launch hypothesis-driven basic and translational investigations about the differential impacts and therapeutic consequences of targeting active mutant KRAS.

We detected no binding or inhibitory effects of our compounds towards the KRAS WT or KRAS G12C proteins, while our data shows that our KRAS G12D inhibitors can efficiently disrupt the KRAS-CRAF interaction. In diverse cancer cells, these molecules can also disrupt activation of MAPK and PI3K/mTOR and can confer anti-proliferative and anti-tumor effects. However, despite this apparently clear picture from our *in vitro* work, we stress that this efficacy is not fully dependent on KRAS mutation status: our data from assays with diverse cancer cells bearing KRAS WT revealed a more complex interaction pattern. Our experiments with xenograft model pancreatic tumors showcase the very promising efficacy and combination therapy synergistic potential of TH-Z835; nevertheless, our observation of weight loss again suggested off-target impacts. A very likely explanation is that the inhibitors bind to and inhibit Rho, Ran, Arf, and/or Rab subfamily proteins that also function in regulating cancer cell proliferation and MAPK signaling. In sum, our study has demonstrated proof-of-concept for a salt-bridge induced-fit pocket strategy to achieve KRAS G12D inhibition, which warrants future medicinal chemistry efforts to ensure specificity and optimal efficacy.

---

**Table S1.** Crystallization data collection and refinement statistics.

|  | KRAS G12D<br>TH-Z816 | KRAS G12D<br>TH-Z827 | KRAS G12D<br>TH-Z835 |
| --- | --- | --- | --- |
| PDB code | 7EW9 | 7EWA | 7EWB |
| **Data collection** |  |  |  |
| Space group | *P*12<sub>1</sub>1 | *P*12<sub>1</sub>1 | *P*12<sub>1</sub>1 |
| Cell dimensions |  |  |  |
| a, b, c (Å) | 47.0, 103.4, 56.3 | 46.7, 102.8, 56.4 | 46.6, 102.5, 56.1 |
| α, β, γ (°) | 90, 106.4, 90 | 90, 107.1, 90 | 90, 106.9, 90 |
| Resolution (Å) | 50.0 – 2.13<br>(2.16 – 2.13) | 50.0 – 2.25<br>(2.28 – 2.25) | 50.0 – 1.99<br>(2.02 – 1.99) |
| No. of observed reflections | 28858 (1177) | 24193 (946) | 34543 (1505) |
| Redundancy | 7.0 (4.0) | 9.1 (6.7) | 6.1 (3.5) |
| Completeness (%) | 99.9 (98.5) | 100.0 (100.0) | 99.9 (100.0) |
| I/sigma (I) | 14.1 (2.4) | 12.6 (3.3) | 10.0 (2.1) |
| *R*<sub>merge</sub> (%)<sup>b</sup> | 7.2 (42.0) | 8.4 (39.8) | 8.8 (39.5) |
| CC<sub>1/2</sub> | 1.0 (0.81) | 1.0 (0.95) | 1.0 (0.88) |
| **Refinement<sup>c</sup>** |  |  |  |
| *R*<sub>work</sub> | 20.1 | 20.3 | 19.5 |
| *R*<sub>free</sub> | 25.7 | 25.8 | 24.6 |
| r.m.s.d bonds (Å) | 0.008 | 0.008 | 0.007 |
| r.m.s.d angles (°) | 1.098 | 1.079 | 1.105 |
| **Ramachandran statistics** |  |  |  |
| Preferred (%) | 96.8 | 97.3 | 98.0 |
| Allowed (%) | 3.0 | 2.7 | 2.0 |
| Outliers (%) | 0.2 | 0 | 0 |
| **Average B-factor (Å<sup>2</sup>) / Atoms** |  |  |  |
| Protein | 39.1 / 4015 | 43.6 / 3902 | 31.1 / 4053 |
| Mg | 36.5 / 3 | 42.0 / 3 | 30.8 / 3 |
| GMPPNP or GDP | 31.8 / 84 | 37.9 / 92 | 26.7 / 92 |
| Inhibitor | 40.1 / 72 | 43.4 / 74 | 32.4 / 111 |
| Solvent | 41.3 / 276 | 43.5 / 169 | 34.4 / 317 |

<sup>a</sup> Values in parentheses are for the highest resolution shell.
<sup>b</sup> $R_{\text{merge}} = \frac{\sum_{hkl} \sum_i |I_i(hkl) - \langle I(hkl) \rangle|}{\sum_{hkl} \sum_i I_i(hkl)}$ , where the sum is over all the *i* measured reflections with equivalent miller indices *hkl*; $\langle I(hkl) \rangle$ is the averaged intensity of these *i* reflections, and the grand sum is over all measured reflections in the data set.
<sup>c</sup> All positive reflections were used in the refinement.

**Fig. S1:**
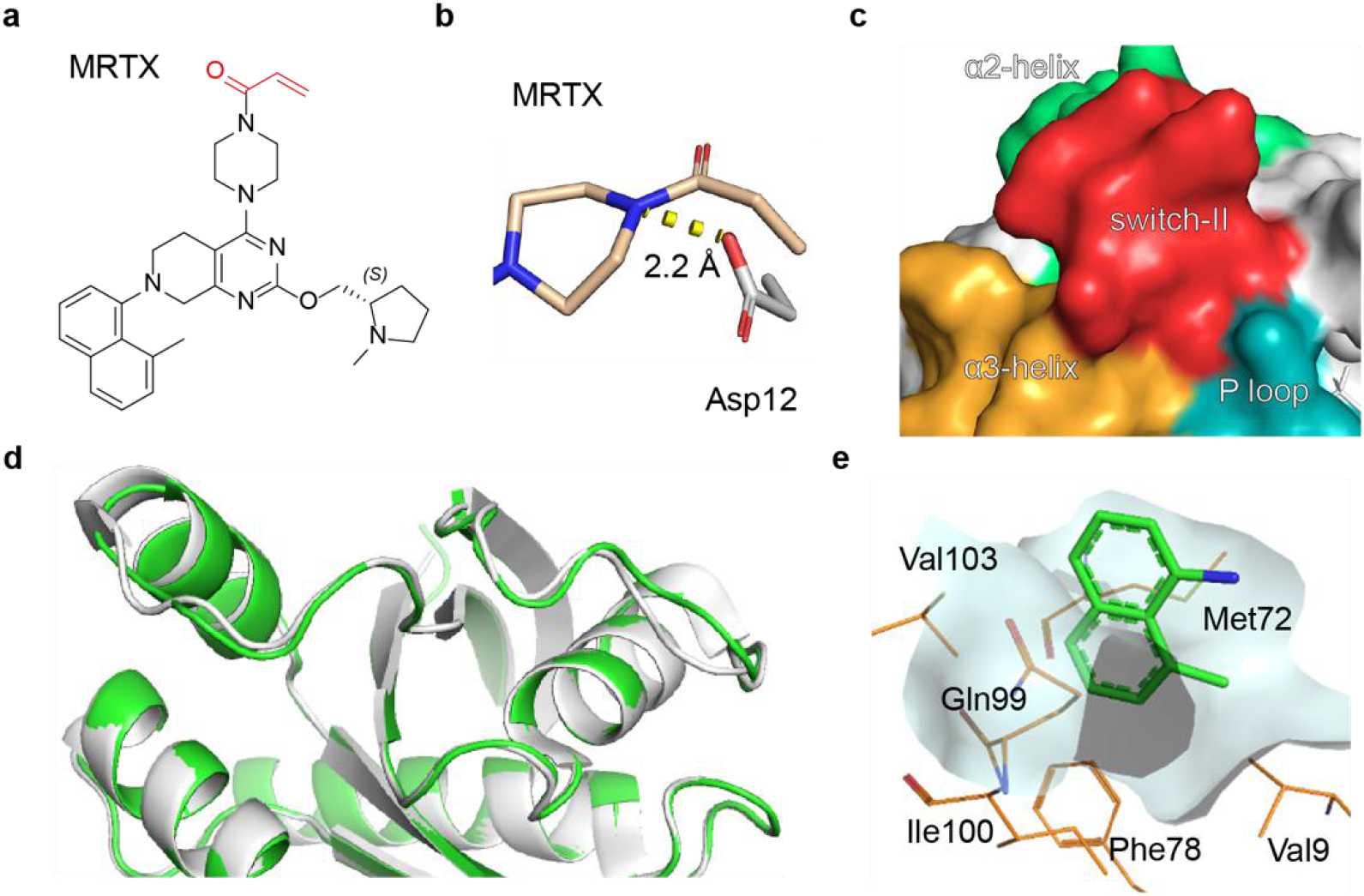
Additional insights into compound binding and the induced-fit pocket of KRAS G12D. **a**, Chemical structure of the previously reported acryloyl-moiety-containing KRAS G12C inhibitor MRTX (the acryloyl moiety is highlighted in red). **b**, Docking pose showing the 2.2 Å distance between the N atom of piperazine moiety (PDB ID 6USX) and the O atom of Asp12 (PDB ID 4EPR). **c**, The inhibitor-free KRAS G12D structure (PDB ID 4EPR). Secondary structures, including α2-helix (green), switch II (red), α3-helix (orange) and P loop (teal), are shown as a surface diagram. **d**, Alignment of TH-Z816-bound KRAS G12D (green, PDB ID 7EW9) and MRTX-bound KRAS G12C (white, PDB ID 6USX) shown as secondary structures. **e**, There is a hydrophobic pocket around the naphthyl moiety of TH-Z816, comprising Val9, Met72, Phe78, Gln99, Ile100, and Val103.

**Fig. S2:**
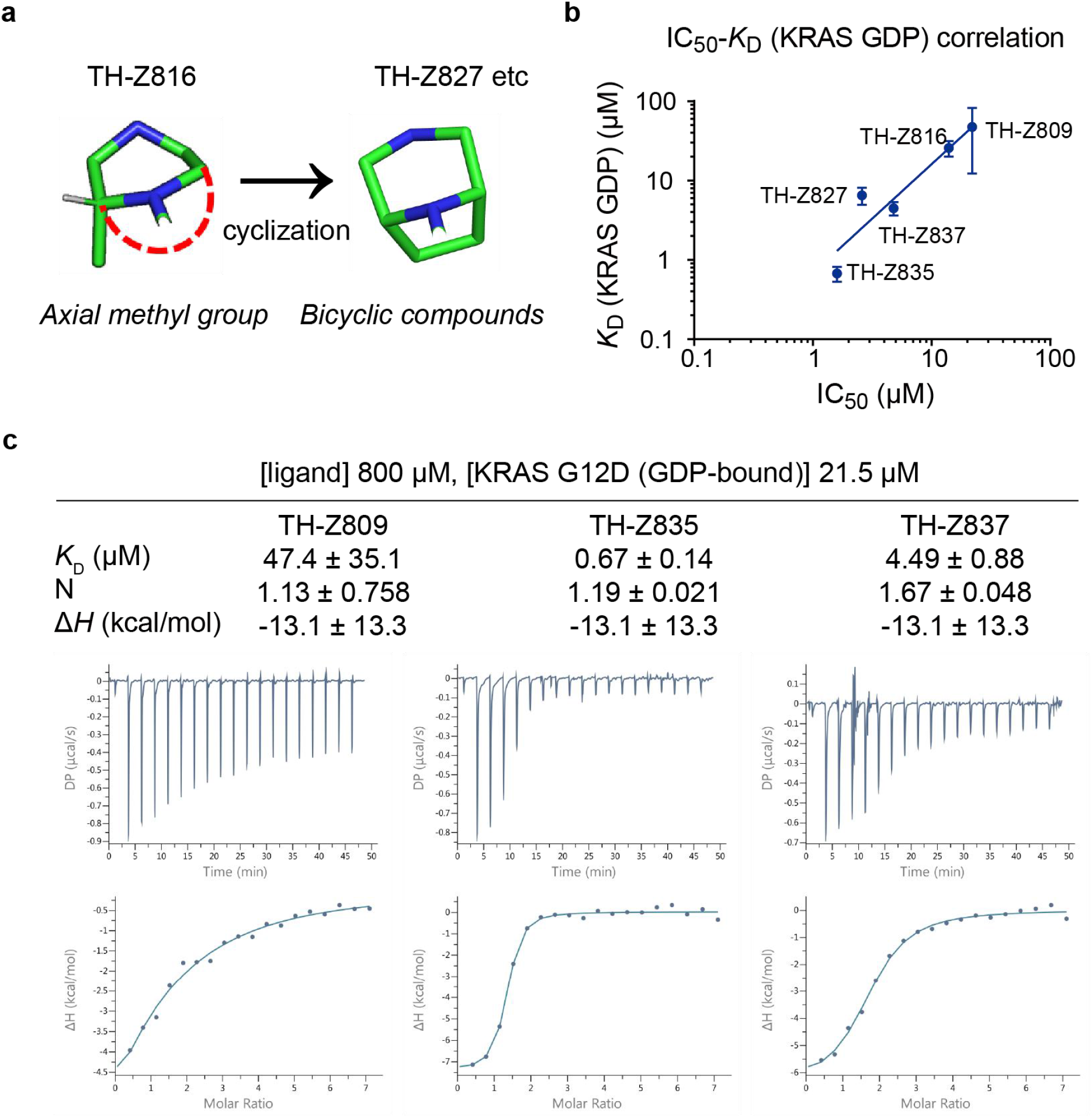
Structural analysis of KRAS G12D bound to TH-Z816. **a**, The axial position of the methyl group of TH-Z816 suggests cyclization as a feasible strategy for inhibitor design. **b**, ITC K_D_ (GDP-bound KRAS G12D) and IC_50_ values of the indicated compounds, fit based on linear correlation (blue line). **c**, ITC assays of the indicated compounds (800 μM) and GDP-bound KRAS G12D (21.5 μM).

**Fig. S3:**
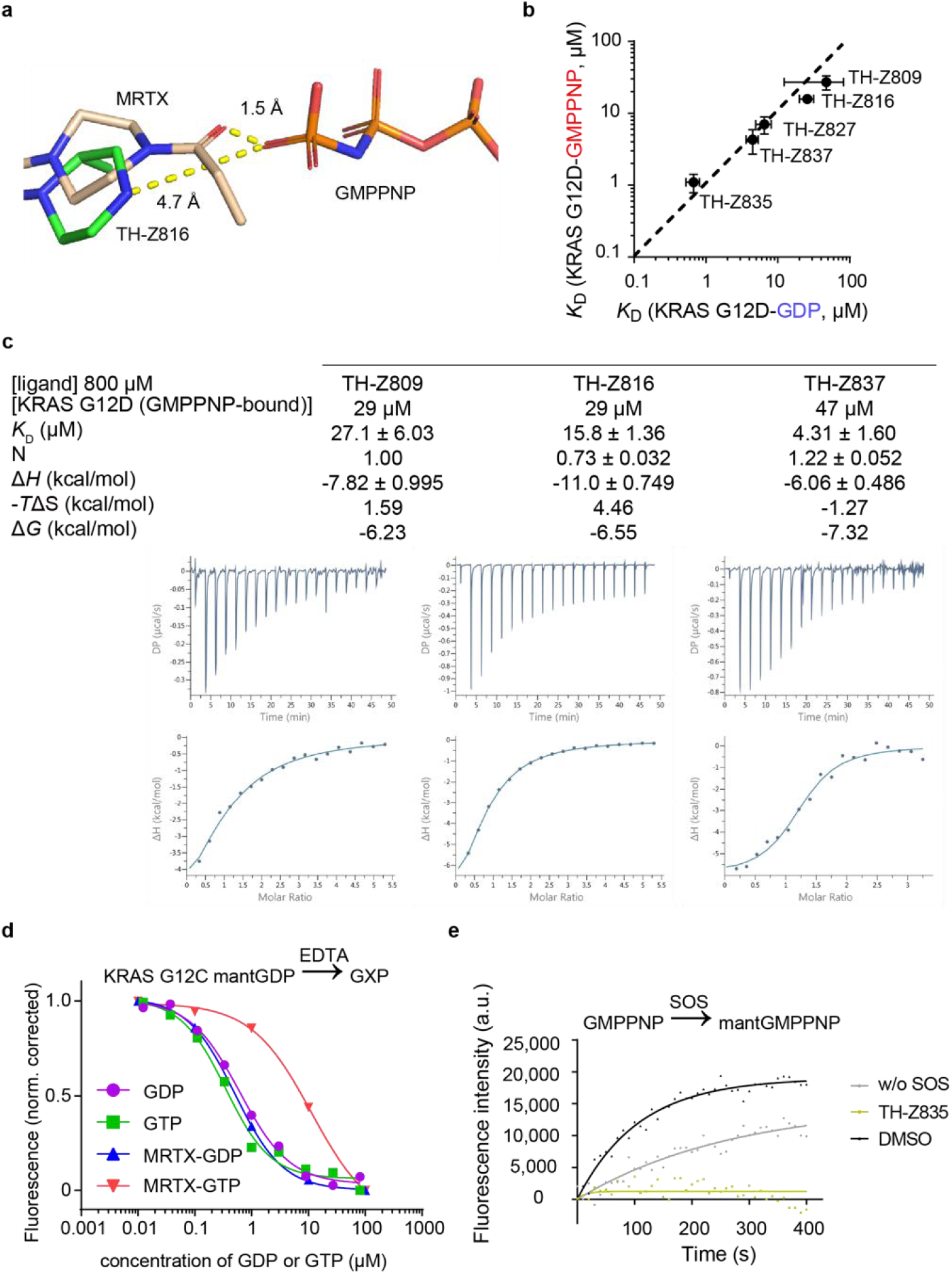
Further analysis of the binding mode of GMPPNP-bound KRAS. **a**, Computational modeling indicates our G12D inhibitor (PDB ID 7EW9) does not have steric clash with the γ-phosphate of GMPPNP (PDB ID 5USJ). The distance between the γ-phosphate of GMPPNP (PDB ID 5USJ) and the acryloyl moiety of the G12C inhibitor MRTX (PDB ID 6USX) is 1.5 Å. The distance between the γ-phosphate of GMPPNP and the piperazine moiety of the G12D inhibitor TH-Z816 (PDB ID 7EW9) is 4.7 Å. The protein structure was modeled using the Protein Preparation Wizard of Schrödinger Maestro. **b**, ITC K_D_ values for each compound for both GMPPNP-bound and GDP-bound KRAS G12D. The dashed line is a linear fitting line (y = x). **c**, ITC assay of the TH-Z809, TH-Z816, or TH-Z837 (800 μM) with GMPPNP-bound KRAS G12D. **d**, EDTA-mediated competition between fluorescently labeled mantGDP loaded on KRAS and free nucleotide (GDP or GTP). The experiment was carried out with KRAS G12C alone (1 μM) or with KRAS G12C treated with MRTX (3 μM). **e**, Inhibitory activity of TH-Z835 measured by SOS-catalyzed nucleotide exchange assays with mantGMPPNP as the incoming nucleotide.

**Fig. S4:**
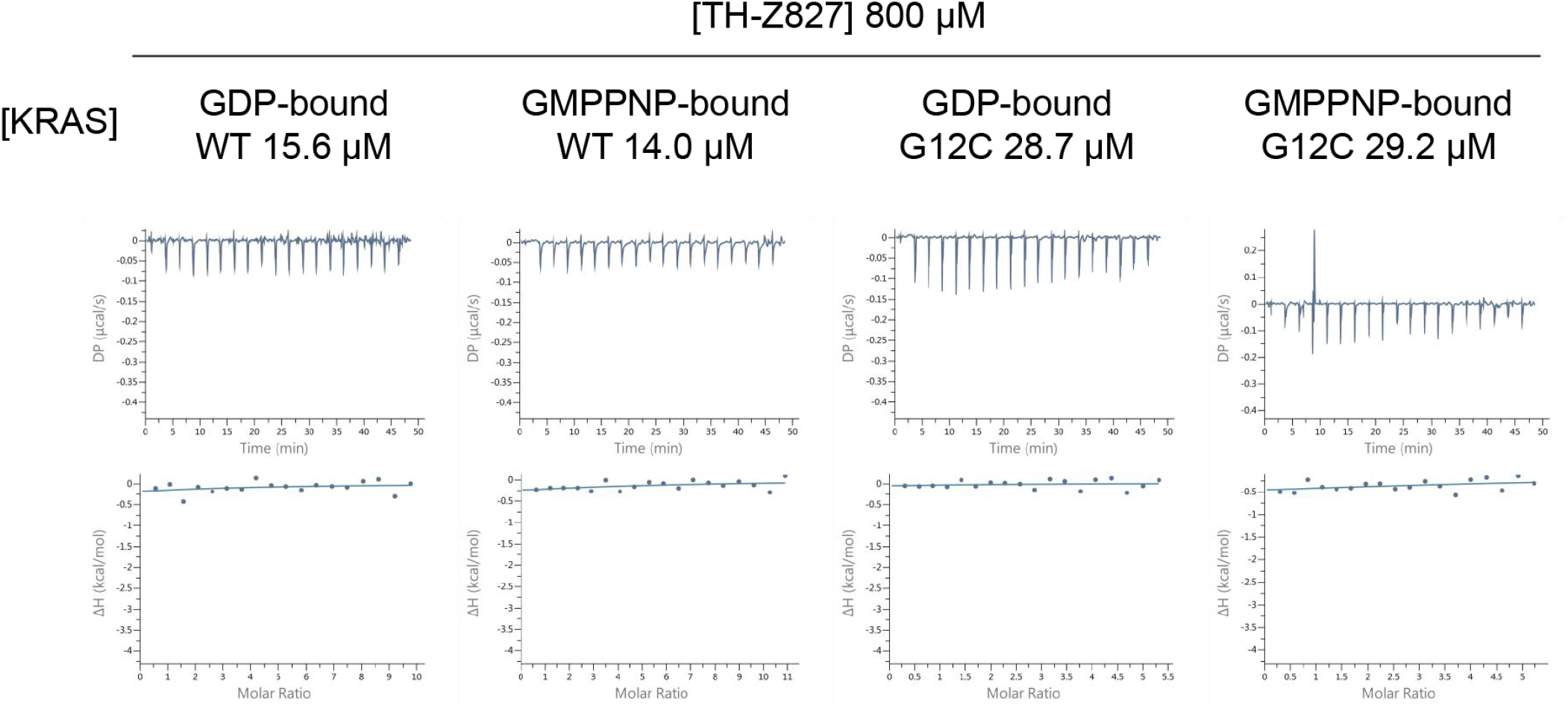
Binding assay of TH-Z827 with GDP- or GMPPNP-bound KRAS (WT or G12C). ITC assays of TH-Z827 (800 μM) and GDP- or GMPPNP-bound KRAS (WT or G12C).

**Fig. S5:**
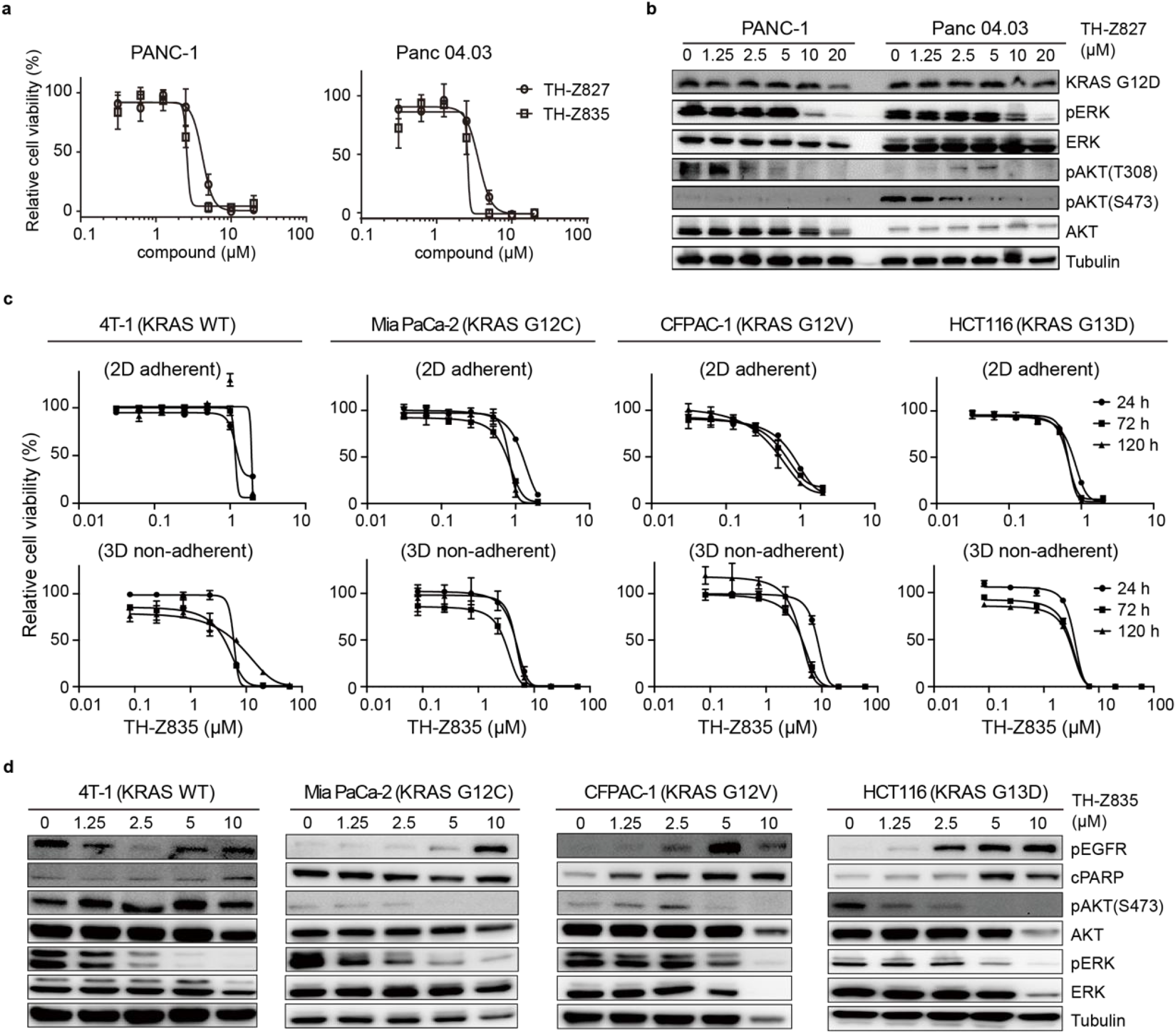
Anti-proliferative effects and signaling inhibition of KRAS G12D inhibitors. **a**, Viability assays of PANC-1 and Panc 04.03 cells treated with indicated concentration of TH-Z827 or TH-Z835 for 24 h. **b**, Immunoblotting analysis of ERK and AKT phosphorylation status in PANC-1 (**left panel**) and Panc 04.03 cells (**right panel**) treated with TH-Z827 for 3 h. **c,** Cell viability assays of 4T1, MIA PaCa-2, CFPAC-1, and HCT116 cells with indicated concentration of TH-Z835 for 24 h, 72h, or 120 h in 2D adherent assays (**upper panel**) and 3D non-adherent assays plates (**lower panel**). **d**, Immunoblotting of EGFR, ERK, AKT phosphorylation, and cPARP in 4T1, MIA PaCa-2, CFPAC-1 and HCT116 cells treated with TH-Z835 for 3 h.

**Fig. S6:**
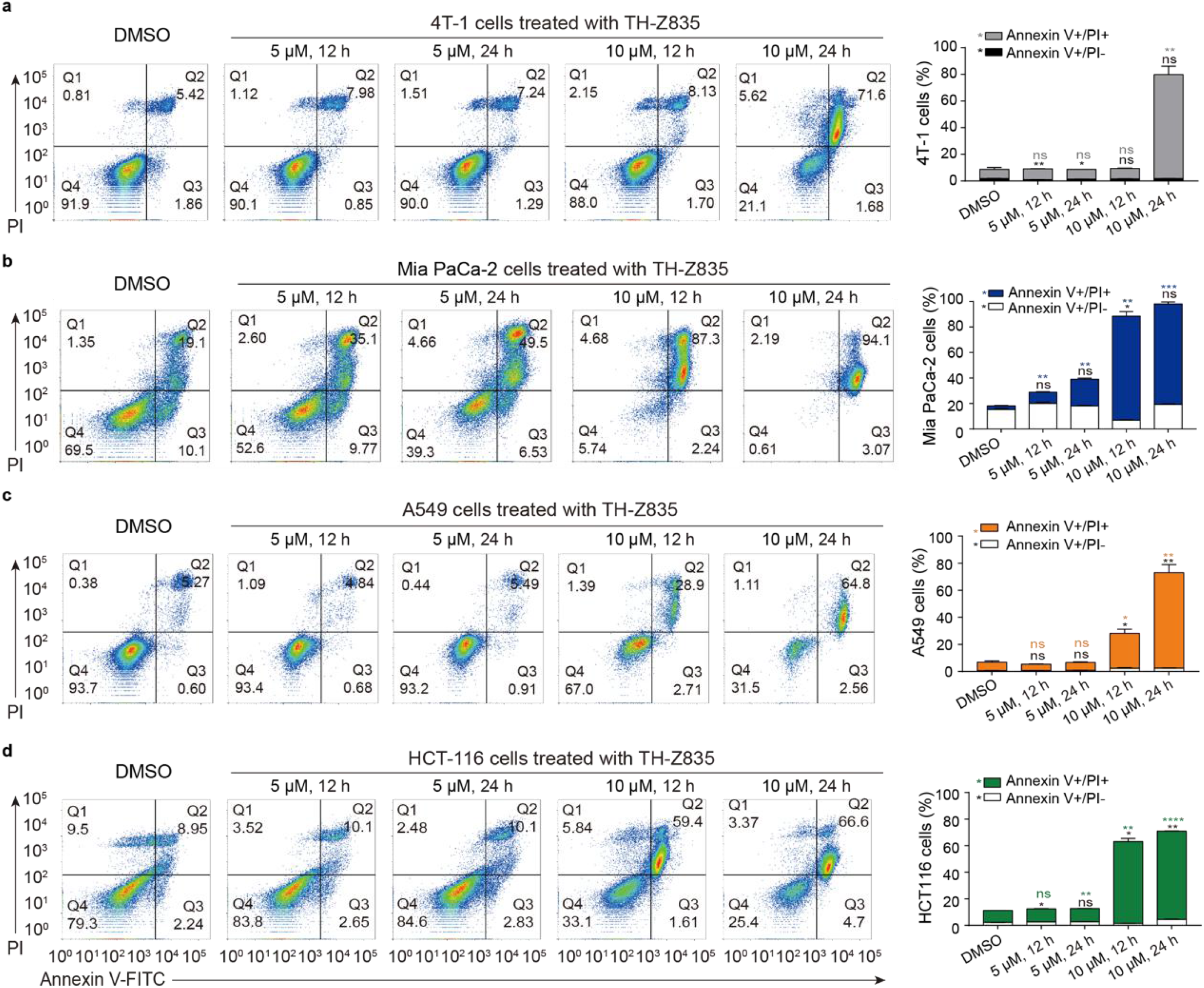
TH-Z835 induces apoptosis in KRAS mutant cells. (**left panel**) Apoptosis analysis by flow cytometry of 4T1 (**a**), MIA PaCa-2 (**b**), A549 (**c**), and HCT116 (**d**) cells upon a 12 or 24-h treatment of TH-Z835. **(right panel)** Apoptotic (Annexin V-positive) cell proportions were quantified. Data are shown as the mean ± s.e.m. (*n* = 3). Student’s t test, * *p* < 0.05, ** *p* < 0.01, *** *p* < 0.001, **** *p* < 0.0001.

**Fig. S7:**
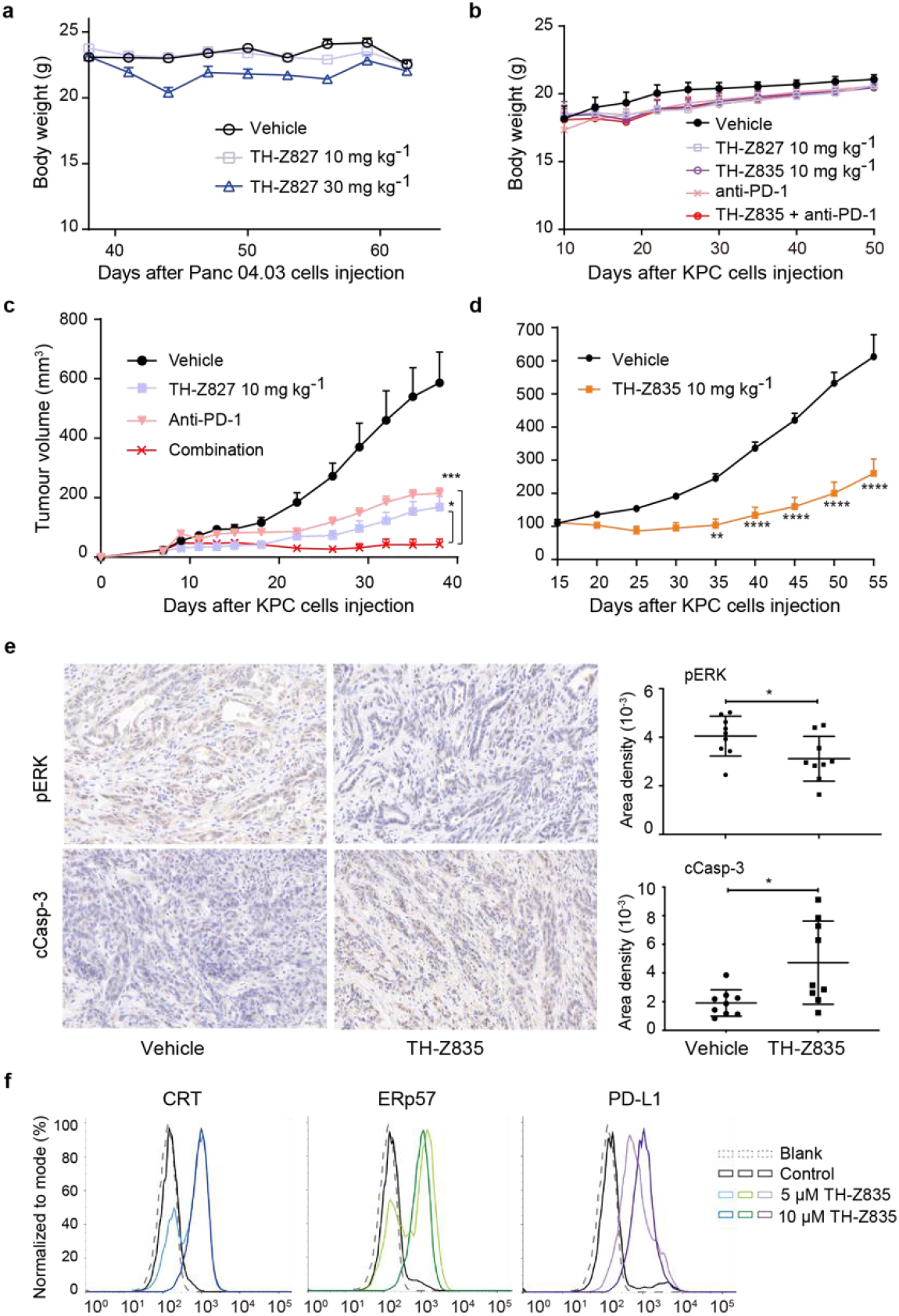
Anti-tumor effects of the KRAS G12D inhibitors alone and in combination with anti-PD-L1 antibody. **a,** Body weight (mean ± s.e.m., *n* = 10) of mice bearing xenograft tumors (from inoculation of Panc 04.03 cells) treated (IP) with TH-Z827 at 10 mg kg^−1^ or 30 mg kg^−1^. b, Body weight (mean ± s.e.m., *n* = 5) of tumour-bearing mice treated (IP) with TH-Z827, TH-Z835, anti-PD-1 antibody or a combination therapy (10 mg kg^−1^ TH-Z835 and 100 μg per dose anti-PD-1 antibody). c, C57BL/6 mice were injected with KPC cells at Day 0, after which TH-Z827, anti-PD-1 antibody, or a combination therapy (10 mg kg-1 TH-Z827 and 100 μg per dose anti-PD-1 antibody) was IP administered according to the indicated dosage schedule shown in Fig. 7b. Combination treatment (*n* = 5, shown as the mean ± s.e.m.) led to a statistically significant decrease in tumour volumes at day 38 compared with either single-agent treatment (one-way ANOVA followed by Dunnett’s test; * P_adj_ < 0.05, *** P_adj_ < 0.001). d, Mice were injected with KPC cells at Day 0 and TH-Z835 at Day 10. The tumour volumes (mean ± s.e.m., *n* =10) were assessed using Student’s t test, ** *p* < 0.01, **** *p* < 0.0001. e, (left panel) Immunohistochemical (IHC) analysis of pERK and cleaved Caspase-3 in tumor section. Scale bar, 20 μm. (right panel) Quantification of IHC positive staining (mean ± s.d., *n* = 9) were assessed using Student’s t test, * *p* < 0.05. f, Flow cytometry analysis of ICD markers (CRT and ERp57) on the surface of KPC cells after 24-h treatment with TH-Z835.

## Methods

### Protein expression and purification

The gene encoding KRAS G12D (residues 1–169) was chemically synthesized and cloned into a pET28a vector with *NdeI* and *XhoI* restriction sites. A construct for the recombinant KRAS 4b G12D was transformed into *E. coli* BL21 (DE3). After the bacterial growth to an OD600 of 0.6, induction was carried out using 1 mM isopropyl-β-D-thiogalactopyranoside (IPTG) (16 °C for 24 h). The cells were lysed and recombinant KRAS G12D was purified using a HisTrap HP (GE Healthcare, 29-0510-21) column. The hexahistidine tag was then removed by TEV-protease and crude protein was further purified. The mature protein was concentrated to 3.5 mg mL^−1^, and the molecular mass of protein was determined by SDS-PAGE (~21 kDa). The KRAS protein sequence is ‘MTEYKLVVVG ADGVGKSALT IQLIQNHFVD EYDPTIEDSY RKQVVIDGET CLLDILDTAG QEEYSAMRDQ YMRTGEGFLC VFAINNTKSF EDIHHYREQI KRVKDSEDVP MVLVGNKCDL PSRTVDTKQA QDLARSYGIP FIETSAKTRQ GVDDAFYTLV REIRKHKEK’. SOS, KRAS WT, and KRAS G12C were expressed and purified in a similar way as KRAS G12D. The plasmid of SOS was provided by Professor Niu Huang of National Institute of Building Sciences (NIBS).

### X-ray crystallography

The protein was further concentrated to 40 mg ml^−1^ for the X-ray crystallography study. For TH-Z816, KRAS G12D protein was directly used for the following procedure. For TH-Z827 and TH-Z835, the endogenous GDP of KRAS G12D was exchanged with GMPPNP catalyzed by EDTA. Next, using the vapor-diffusion method, thin plates were observed after a week at 20 °C under the crystallization conditions of 0.2 M sodium acetate, 0.1 M Tris, pH 8.5, 26 % (w/v) PEG 3350. To obtain the complex structures, the protein crystals were soaked into 2 mM inhibitor for 6 h. The soaked crystals were cryoprotected in the mother liquor supplemented with 10% glycerol prior to flash-freezing. The X-ray diffraction data was obtained at the in-house beamline BRUKER D8 VENTURE at Hubei University. Datasets were initially processed with PROTEUM3 v2020.6, solved by molecular replacement using Phaser with KRAS (PDB ID 4EPR), and refined to the indicated statistics using PHENIX 1.10.1-2155 and Coot 0.8.3^41^. The figures were drawn using PyMOL.

### EDTA-catalyzed nucleotide exchange

Endogenous nucleotides in KRAS were exchanged with GDP (Sigma, G7127), GMPPNP (Sigma, G0635), mantGDP (Jena Bioscience, NU-204S), or mantGMPPNP (Jena Bioscience. NU-207S) using a previously reported EDTA-catalyzed procedure^42,43^. Briefly, KRAS G12D protein (10 μM) was incubated with incoming nucleotide (200 μM) and EDTA (2.5 M) for 1.5 hr at room temperature. After incubation, the sample was put on ice for 2 min, and then MgCl_2_ (5 mM final) was added to stop the reaction. Excess unbound nucleotide was removed using a NAP-5 column (GE Healthcare, 17085302).

### SOS-catalyzed nucleotide exchange

For studies with mant-nucleotide loaded protein, 12 μL of the protein (1.25 μmol L^−1^) in reaction buffer containing 1 mmol L^−1^ of a given incoming nucleotide (GDP or GMPPNP) was added to a 96-well half-area microplate (Corning 3686). After incubation with compounds for 10 min, reactions were initiated by the addition of 3 μL of SOS (10 μmol L^−1^); fluorescence (λ_ex_ = 355 nm, λ_em_ = 460 nm) was monitored for 45 minutes at 60-second intervals with a multimode microplate reader (PerkinElmer, EnVision). For the exchange assays with incoming mant-nucleotide, 12 μL of the purified protein (1.25 μmol L^−1^) was used (in reaction buffer containing 1 μmol L^−1^ of the indicated incoming nucleotide). Fluorescence data were fitted to a one phase decay model with GraphPad Prism 7.0.

### Molecular docking

Compounds were constructed using the Schrödinger Maestro 3D sketcher, and candidate conformations were prepared and generated using LigPrep. Protein structures were optimized using Protein Preparation Wizard software, with docking grids profiles generated around the ligand. Docking was performed using Glide software, with standard precision. The force field was OPLS_2005.

### ITC assays

ITC experiments were carried out on at 25 °C with 19 injections, 2 μL per injection, and 150 s intervals using a MicroCal PEAQ-ITC instrument (GE Healthcare). Buffer was exchanged into 25 mM Tris-HCl, 100 mM NaCl, 5 mM MgCl_2_, 1 mM TCEP (Sigma, C4706), 0.05 % Tween-20 (Sigma, P1379) and 2.5 % (v/v) DMSO. The protein was loaded into a cell, and the compound was titrated. Reference power was set to 10 μcal sec^−1^, and the compound solution was titrated at 150 s. Data were fitted into a one site model, and KD, N, ΔG, ΔH, and −TΔS values were calculated using MicroCal Analysis software.

### KRAS-CRAF interaction assay

Three KRAS mutant cell lines were constructed based on HEK 293T cells using plasmids including KRAS WT-CRAF, KRAS G12C-CRAF, and KRAS G12D-CRAF. Cells were cultured on 6-well plates (2.5×10^5^ per well) and treated with 1 μg ml^−1^ doxycycline for 24 h. Then each well was treated with 200 μL lysis buffer (Promega, E2661) and centrifuged at 4 °C for 5 min. The supernatant was collected and incubated with compounds (TH-Z827, MRTX, or BI-2852) for 10 min. Then 20 μL luciferin substrate buffer (Promega, E2510) was added and incubated for 10 min before luminescence was measured with a multimode microplate reader (PerkinElmer, EnVision). Data were fitted into an inhibitor-response model to get IC_50_ values.

### 2D cell viability assays

All the cell lines used in this study were obtained from ATCC. PANC-1 and Panc 04.03 cells were seeded onto 96-well microplates (5×10^3^ cells per well) and cultured for 24 h with DMEM medium (Gibco, 11960-051) supplemented with 10% FBS (Biological Industries, 04-001-1) and 1% Pen/Strep (Beyotime, C0222). After treatment with compounds (TH-Z827, MRTX, or BI-2852), cells were incubated for another 24 h. Cell Counting Kit-8 reagents (Beyotime, C0042) were added and incubated for another 1 h. Cell viability was measured at OD 450 nm using a multimode microplate reader (PerkinElmer, EnVision). Data were fitted into an inhibitor-response model to get IC_50_ values.

### 3D cell viability assays

For comparison of anti-growth activity, a CellTiter-Glo (CTG) 3D cell viability assay (Promega, G9682) was used. Cells (1×10^4^ cells per well) were seeded (using the same media) in ultra-low attachment surface 96-well format plates (Corning Costar #3474). The day after plating, cells were treated with a 7 point 3-fold dilution series of indicated compounds (200 ul final volume per well) and cell viability was monitored at 1, 3, 5 days according to the manufacturer’s recommended instructions, where 50 ul of CellTiter-Glo reagent was added, vigorously mixed, covered, and placed on a plate shaker for 20 min to ensure complete cell lysis prior to assessment of luminescent signal.

### Colony formation assay

KPC or PANC1 (1×10^3^ cells per well) were seeded in the 6-well plates in triplicate. After overnight incubation, cells were treated with various concentrations of TH-Z835 or vehicle (DMSO), and allowed to grow for 10 to 14 days, during which medium was changed every 3 days. After 10 to 14 days, cells were then fixed by 4% paraformaldehyde (Leagene, DF0135) followed by adding crystal violet solution (Beyotime, C0121) for 30 min. After washing with water, pictures of the wells were taken by microscope. Then the stain was extracted by adding 10% methanol-acetic acid solution, and the absorbance was measured at 590 nm.

### Cell cycle detection by flow cytometry

Cells were seeded in 6-well plate and synchronized with serum-free medium for 24 h. Next, the cells were released in complete medium containing either DMSO or TH-Z835, and collected for analysis at 24h. For cell cycle analysis, the cell DNA was stained with propidium iodide (PI) using cell cycle and apoptosis analysis kit (Beyotime, C1052). Briefly, cells were harvested by trypsinization and fixed with cold 75 % ethanol at 4 °C overnight. The fixed cells were collected and suspended in PBS containing 10 μg ml^−1^ PI and 10 μg ml^−1^ RNase A, and then incubated at room temperature for 30 min. DNA content was analyzed by the BD FACS Calibur (BD Biosciences), and each histogram was constructed with the data from at 10,000-20,000 events. The data were analyzed and expressed as percentages of total gated cells using the Modfit LT™ Software (BD Biosciences).

### Western blotting

PANC-1 and Panc 04.03 cells were cultured on 6-well plates (5×10^5^ cells per well) and treated with TH-Z827 for 3 h. Protein samples were prepared, electrophoresed (10% SDS-PAGE), and transferred to a polyvinylidene fluoride (PVDF) membrane. RAS G12D (Cell Signaling Technology, 14429), pERK1/2 (Cell Signaling Technology, 4370), ERK1/2 (Cell Signaling Technology, 4695), pAKT(Thr308) (Cell Signaling Technology, 2965), pAKT(Ser473) (Cell Signaling Technology, 4060), AKT (Cell Signaling Technology, 2920) and tubulin (Proteintech, 66240-1-Ig), EGF Receptor (Cell Signaling Technology, 4267), phospho-EGF Receptor (Tyr1068) (Cell Signaling Technology, 3777), cleaved PARP (Asp214) (Cell Signaling Technology, 9541), cleaved Caspase-3 (Asp175) (Cell Signaling Technology, 9661), cleaved Caspase-7 (Asp198) (Cell Signaling Technology, 9491), cell cycle regulation antibody sampler kit (Cell Signaling Technology, 9932), antibodies were used to detect the individual proteins. For MAPK signaling in KRAS G12D or non-G12D mutant cells, cells were cultured on 6-well plates (5×10^5^ cells per well) and treated with a serial titration of TH-Z835 or TH-Z827 for 3 h. For apoptosis, RTK feedback regulation, or cell-cycle arrest assay, PANC-1 or KPC were cultured on 6-well plates (5×10^5^ cells per well) and treated with a serial titration of TH-Z835 at time points for up to 72 h.

### Apoptosis detection by Annexin V binding

Cells (5×10^5^ cells per well) were seeded in 6-well plate overnight. Next, the cells were treated with either DMSO or TH-Z835 for 12 h or 24 h. For apoptosis analysis, cells were harvested by trypsinization and washed twice with ice-cold PBS, then stained with Annexin V-FITC and PI by Apoptosis Detection Kit (Beyotime, C1062) in the dark at room temperature for 10 min. Then cells were analyzed with the BD FACS AriaII and FlowJo software.

### Apoptosis detected by WB

For apoptosis, RTK feedback regulation, or cell-cycle arrest assay, PANC-1 or KPC were cultured on 6-well plates (5×10^5^ cells per well) and treated with a serial titration of TH-Z835 at various time points for up to 72 h.

### Immunogenic cell death and PD-L1 detection by flow cytometry

The cells were treated with TH-Z835 as indicated for 24 h before harvesting. After washing twice in cold PBS, cells were incubated for 30 min with anti-PD-L1 (ab213480, Abcam), anti-CRT (1:1000, ab2907, Abcam), or anti-ERp57 (1:1000, ab10287, Abcam) antibody, diluted in cold blocking buffer (2% FBS in PBS), followed by washing and incubation with the Alexa Fluor 488-labeled secondary antibody (1:1000, ZF-0511, ZSGB-BIO) for 30 min. Then cells were analyzed with the BD FACS AriaII and FlowJo software.

### Real-Time quantitative PCR

PANC-1 were treated with multiple doses of TH-Z835 for 24 h. Total RNA was extracted with TRIzol reagent (CWBIO, CW0580). cDNA was prepared using 1 μg of RNA with a cDNA Synthesis Kit (Yeasen, 11141ES10). SYBR-green-based qPCR was performed using primers for PD-L1 (forward, TTTGCTGAACGCCCCATACA; reverse, TTGGTGGTGGTGGTCTTACC), GAPDH (forward, GAGTCAACGGATTTGGTCGT; reverse, TTGATTTTGGAGGGATCTCG). Gene expression was calculated by the comparative ΔΔCT method with the GAPDH for normalization.

### Mouse models

BALB/c nude mice were subcutaneously injected with Panc 04.03 cells (1×10^7^ per dose) at Day 0. Mice were randomized when the mean tumour volume was ~70 mm^3^. Each group of mice (*n* = 10) were IP injected with PBS, 10 mg kg^−1^ or 30 mg kg^−1^ TH-Z827) according to the dosage schedule from Day 38 to Day 62.

C57BL/6 mice were subcutaneously injected with KPC (KrasLSL.G12D/+; p53R172H/+; PdxCretg/+) cells (5×10^5^ per dose). Mice were randomized when the mean tumour volume was ~20 mm^3^. Mice of each group (*n* = 10) were IP injected with PBS, anti-PD-1 antibody (100 μg per dose, Bio X Cell, BE0033-2), 10 mg kg^−1^ TH-Z827, or a combination (10 mg kg^−1^ TH-Z827 and anti-PD-1 antibody) according to a pre-defined dosage schedule from Day 7 to Day 38. Polyclonal Armenian hamster IgG (Bio X Cell, BE0091) was used as a control antibody.

Statistical analysis of differences in mean tumour volume between vehicle and treated groups were assessed using a one-way ANOVA test conducted in GraphPad Prism. A *P* value < 0.05 was considered statistically significant.

### Immunohistochemistry and immunofluorescence

Tumor samples were obtained after treatment for 30 days, and fixed in 4.0% paraformaldehyde solution (Leagene, DF0135), embedded in paraffin, and cut into 4μm sections. Sections were used for Immunohistochemistry (IHC) and Immunohistochemistry (IF) according to the standard procedures. The IHC staining protocol was briefly described as follows: the slides were routinely deparaffinized, rehydrated, subjected to antigen retrieval, and incubated in 3% hydrogen peroxide to block endogenous peroxidase. Subsequently, the slides were blocked with 3% BSA and incubated with primary antibodies against pERK1/2 (1:500, Cell Signaling Technology, 4370) or cleaved Caspase-3 (1:400, Cell Signaling Technology, 9661) at 4 °C overnight, and with polymer-HRP-conjugated anti-rabbit secondary antibody (1:200S, Servicebio, GB23303). Then, the sections were stained with DAB kit (Servicebio, G1211), counterstained with hematoxylin, dehydrated, and cover-slipped. For IF staining, slides were incubated with primary antibodies against pERK1/2 (1:200, Cell Signaling Technology, 4370), cleaved Caspase-3 (1:200, Cell Signaling Technology, 9661) at 4 °C overnight, and with CY3-conjugated anti-rabbit secondary antibody (1:300, Servicebio, GB21303). Then, the sections were counterstained with antifading agent/DAPI (Beyotime, P0131).

## Data availability

Most of the data generated or analyzed during this study are included in this Article or available as supplementary information. X-ray crystallographic coordinates and structure factor files have been deposited in the Protein Data Bank (PDB IDs 7EW9, 7EWA, 7EWB). Other data that support the findings of this study are available from the corresponding authors.

## Acknowledgements

We acknowledge Professor Niu Huang for providing the plasmid of SOS and Dr. Yifeng Xia for donating the HEK 293T cell lines used for detecting KRAS-CRAF interactions. This work was supported by the National Key Research and Development Program of China (2021YFC2100300), the Beijing Natural Science Foundation (Z190015), the National Natural Science Foundation of China (81991492), the Beijing Advanced Innovation Centre for Structural Biology, the Beijing Advanced Innovation Centre for Human Brain Protection and Tsinghua University Spring Breeze Fund.

## Author contributions

Y.Z. initiated, designed and supervised the project, and wrote the manuscript. Z.M. designed, synthesized compounds, performed enzymatic assays, and wrote the manuscript. H. X., Y-Y. Z and Y. D performed the cellular experiments. H. X. performed the animal experiments and wrote the manuscript. P.S., Y.Y., J.X., L.Z., and Y.-Y.Y. performed protein purification and structural studies. Y.S., C.-C.C., and R.-T.G. contributed to the study design and discussion.

## Notes

### Competing Interest Statement

The authors have declared no competing interest.

